# Production of Noncapped Genomic RNAs is Critical to Sindbis Virus Disease and Pathogenicity

**DOI:** 10.1101/2020.06.11.145847

**Authors:** Autumn T. LaPointe, V. Douglas Landers, Claire E. Westcott, Kevin J. Sokoloski

## Abstract

Alphaviruses are positive-sense RNA viruses that utilize a 5’ cap structure to facilitate translation of viral proteins and to protect the viral RNA genome. Nonetheless, significant quantities of viral genomic RNAs that lack a canonical 5’ cap structure are produced during alphaviral replication and packaged into viral particles. However, the role/impact of the noncapped genomic RNA (ncgRNA) during alphaviral infection *in vivo* has yet to be characterized. To determine the importance of the ncgRNA *in vivo*, the previously described D355A and N376A nsP1 mutations, which increase or decrease nsP1 capping activity respectively, were incorporated into the neurovirulent AR86 strain of Sindbis virus to enable characterization of the impact of altered capping efficiency in a murine model of infection. Mice infected with the N376A nsP1 mutant exhibited slightly decreased rates of mortality and delayed weight loss and neurological symptoms, although levels of inflammation in the brain were similar to wild type infection. The mice infected with the D355A nsP1 mutant showed significantly reduced mortality and morbidity compared to mice infected with wild type virus. Interestingly, both capping mutants had roughly equivalent viral titer in the brain compared to wild type virus, illustrating that the changes in mortality were not due to deficits in viral replication or dissemination. Examination of the brain tissue revealed that mice infected with the D355A capping mutant had significantly reduced cell death and immune cell infiltration compared to the N376A mutant and wild type virus. Finally, expression of proinflammatory cytokines was found to be significantly decreased in mice infected with the D355A mutant, suggesting that capping efficiency and the production of ncgRNA are vital to eliciting pathogenic levels of inflammation. Collectively, these data indicate that the ncgRNA have important roles during alphaviral infection and suggest a novel mechanism by which noncapped viral RNA aid in viral pathogenesis.

**AUTHOR SUMMARY:** Mosquito transmitted alphaviruses have been the cause of widespread outbreaks of disease which can range from mild illness to lethal encephalitis or severe polyarthritis. In order to successfully replicate, the alphavirus RNA genome needs a 5’ cap structure so that the genome can be translated and produce the viral replication machinery. Despite this, a large number of viral genomes produced during infection do not have a 5’ cap structure, and their role during infection is unknown. Using mouse models of infection and point mutations in the nsP1 protein of Sindbis virus which alter the amount of noncapped genomic RNA (ncgRNA) produced, we found the decreasing the production of ncgRNA greatly reduced morbidity and mortality as well as proinflammatory cytokine expression, resulting in less tissue-damaging inflammation in the brain. These studies suggest that the ncgRNAs contribute to pathogenesis through the sensing of the ncgRNAs during alphaviral infection and are necessary for the development of severe disease.

## INTRODUCTION

Alphaviruses are positive-sense, single-stranded RNA arboviruses that are capable of causing severe disease. The natural enzootic transmission cycle of these viruses is between a mosquito vector and a mammalian host, typically rodents or birds, although epizootic spillover events can occur that result in infection of humans and equids. Alphaviruses are broadly categorized as either arthritogenic or encephalitic based on disease symptomology. The arthritogenic alphaviruses, such as Chikungunya virus (CHIKV) and Ross River Virus (RRV) are capable of causing disease ranging from mild febrile illness to severe polyarthralgia, which can persist anywhere from weeks to years following infection [1–3]. In contrast, the encephalitic alphaviruses, such as Venezuelan Equine Encephalitis virus (VEEV) and some strains of Sindbis virus (SINV), like the AR86 strain used in this study, can cause mild to severe neurological symptoms, including encephalitis that can potentially lead to the death of the host [3–5]. While alphaviruses pose a large threat to public health, there are currently no safe and effective vaccines or antiviral therapies to prevent or treat alphaviral disease.

Alphaviruses produce three RNA species during infection: the genomic strand, which encodes the nonstructural proteins; the minus-strand RNA template; and the subgenomic RNA, which encode the structural proteins. Both the genomic and subgenomic RNAs have a type 0 cap structure added to their 5’ ends to facilitate translation and protect the 5’ end of the transcripts [6–8]. The addition of the cap structure to the 5’ end of viral RNAs is primarily carried out by nonstructural proteins 1 and 2 (nsP1, nsP2). NsP2 removes the 5’ γ-phosphate from the nascent vRNA molecule while, in a separate reaction, the methyltransferase domain of nsP1 catalyzes the addition of a methyl group from S-andenosylmethionine to a GTP molecule, forming a covalent m7GMP-nsP1 intermediate [9, 10]. The m7GMP moiety is then transferred to the 5’end of the vRNA molecule by the guanylyltransferase activities of nsP1, resulting in the 7meGppA type 0 cap structure [11].

In response to the lack of preventatives or treatments, targeting the alphaviral replication machinery has been a popular approach for developing potential antiviral therapies. Capping of the genomic and subgenomic vRNA is vital for successful viral replication, as mutations which completely inhibit capping of the viral RNA render the virus noninfectious. Thus, because nsP1 is responsible for the alphaviral capping process, it has been a popular target for antiviral research. In particular, a number of compounds have been developed which inhibit nsP1 capping activity and reduce viral replication *in vitro*, but, to date, none have been tested for efficacy against alphaviral infection *in vivo* [12–15]. In addition to the development of drugs against nsP1 activity, multiple residues in nsP1 have also been identified as determinants for alphaviral virulence, however the impact of these residues on alphaviral capping efficiency has never been delineated. The SINV nsP1/nsP2 cleavage mutant T538I has been shown to determine pathogenicity in mouse models of infection by altering nonstructural polyprotein processing and the virus’ sensitivity to interferon [16, 17]. More recently, a group of six mutations in the nsP1 of RRV have also been shown to attenuate alphaviral disease in mice, although the mechanism of attenuation and the impacts of these mutations on alphaviral replication have yet to be fully characterized [18, 19]. These studies illustrate the significance of nsP1 to alphaviral infection and pathogenicity, but have yet to determine the importance of alphaviral capping efficiency and the production of the ncgRNAs to *in vivo* infection.

While capping of the viral RNA is critical to viral protein expression and viral replication, we have previously shown that the genomic vRNA are not universally capped, and that a significant proportion of the alphaviral genomic RNA produced and packaged during infection lack the 5’ cap structure [20]. In addition, our recently published study showed that the proportion of noncapped genomic RNA (ncgRNA) produced during SINV infection could be altered using point mutations in nsP1 to modulate capping activity [21]. Specifically, incorporating a D355A mutation in the nsP1 of SINV resulted in increased capping efficiency, and therefore decreased ncgRNA production, relative to wild type SINV. Alternatively, a N376A mutation in nsP1 resulted in decreased capping efficiency and increased ncgRNA production. By utilizing these mutations to alter ncgRNA production, we were able to show that increasing the capping efficiency of nsP1 was detrimental to SINV infection in tissue culture models of infection, while decreasing nsP1 capping efficiency did not significantly affect viral titer or overall replication.

However, the presence or lack of a phenotype *in vitro* is not always indicative of what will occur during infection *in vivo*. As such, the goal of this study was to determine the effect of altered ncgRNA production on alphaviral pathogenesis by using the previously described nsP1 capping mutants in a mouse model of infection. The data presented here show that modulating ncgRNA production through the use of the D355A and N376A point mutations to alter nsP1 capping efficiency in nsP1, has a profound impact on alphaviral pathogenesis. In specific, decreasing capping efficiency resulted in increased sensitivity to type-I IFN and a slight decrease in mortality. Surprisingly, increasing capping efficiency resulted in almost complete abrogation of morbidity and mortality, despite showing increased resistance to type-I IFN, due to reduced immune infiltration and production of inflammatory cytokines in the brain. Collectively, our findings indicate that the ncgRNA are important in determining the host immune response to viral infection and play a critical novel role in alphaviral pathogenesis.

## MATERIALS AND METHODS

### Tissue culture cells

BHK-21 cells (a gift from Dr. R. W. Hardy, Indiana University – Bloomington) and L929 cells (a gift from Dr. P. Danthi, Indiana University - Bloomington) were maintained in minimal essential media (MEM; Cellgro) containing fetal bovine serum (FBS; Atlanta Biologicals), 1% Penicillin-Streptomycin (Cellgro), 1% nonessential amino acids (Cellgro), and 1% L-glutamine (Cellgro). SK-N-BE(2) nerve cells (a gift from Dr. L Beverly, University of Louisville) were maintained in Dulbecco’s Modified Eagle Medium (DMEM)/F12 media containing 10% FBS, 1% Penicillin-Streptomycin, and 1% L-glutamine. All cell lines were cultured at 37° C and 5% CO_2_ in a humidified incubator. Regular passaging using standard subculturing techniques was used to maintain low passage stocks.

### Generation of AR86 SINV capping mutants

The AR86 SINV nsP1 mutants used in this study were generated by Gibson Assembly via the use of a Gibson Assembly HiFi 1-step kit (SGI), using a restriction-digested AR86 cDNA plasmid and a synthetic DNA fragment, according to manufacturer’s instructions [5]. Mutants were verified by whole-genome sequencing; full-genome sequences are available upon request.

### Production of wild-type and mutant SINV Stocks

Wild-type, D355A, and N376A SINV AR86 were prepared by electroporation, as previously described [22]. Approximately 2.8×10^6^ BHK-21 cells were electroporated with 10μg of *in vitro*-transcribed RNA. This was done using a single pulse at 1.5kV, 12mA, and 200 Ω from a Gene Pulse Xcell system (BioRad) as previously described (LaPointe et al, 2018 JVI). Afterwards, cells were incubated under normal conditions until cytopathic effect was apparent, at which point the supernatant was collected, clarified via centrifugation at 10,000xg for 10min at 4°C, and aliquoted into small volume stocks which were stored at −80°C for later use.

### Analysis of viral growth kinetics

To determine if the mutation of the SINV nsP1 protein negatively impacted AR86 SINV infection, one-step viral growth kinetics for each capping mutant were assayed in tissue culture models of infection. BHK-21 cells were seeded in a 12-well plate and incubated under normal conditions until cell monolayers were 80-90% confluent. The cells were then infected with either wild-type virus or the individual capping mutant virus at a multiplicity of infection (MOI) of 5 infectious units (IU) per cell and the virus was allowed to adsorb for 1 h. The inoculum was then removed, the cells were washed with 1x phosphate buffered saline (PBS) to remove any unbound viral particles, and whole medium supplemented with 25mM HEPES was added. The cells were incubated at 37° C and tissue culture supernatants were harvested (and the media replaced) at the indication times post infection. Viral titer was then determined via plaque assay.

### Quantification of infectious virus by plaque assay

In order to determine infectious viral titer of all viral samples produced during this study, standard virological plaque assays were used. To summarize, BHK-21 cells were seeded in 24-well plates under normal incubation conditions until the cell monolayers were 80-90% confluent. At that point, the cells were inoculated with 10-fold serially dilutions of virus containing samples followed by a 1-hour adsorption period. Afterwards, cells were overlaid with a solution of 0.5% Avicel (FMC Corporation) in 1x media for 48 h [23]. The monolayers were then fixed with formaldehyde solution (3.8% formaldehyde-1x PBS) for at least 1 h. The overlay was then removed, and the plaques were visualized via crystal violet staining.

### Type-I IFN sensitivity assay

L929 cells were seeded in a 48 well plate and, upon reaching 80-90% confluency, were inoculated with either wild-type parental virus or one of the individual capping mutants at an MOI of 10 IU/cell. After a 1 h adsorption period, the inoculum was removed, the cells were washed twice with 1x PBS, and whole media was added. At the indicated times post infection, 20 international units (IU) of type-I IFN was added to the media. Supernatants were collected at 24hpi and viral titer was determined by plaque assay.

### Detection of ISG and IFNβ transcripts

To determine abundance of IFNβ and the listed ISG transcripts, L929 cells were seeded in a 24-well plate and, upon reaching 80-90% confluency, were inoculated with wild-type SINV or one of the capping mutants at an MOI of 10 IU/cell. Additionally, L929 cells were mock infected with PBS to determine baseline IFNβ and ISG expression. After a 1h adsorption period, the inoculum was removed, the cells were washed once with 1× PBS, and whole media was added. At the specified timepoints, media was removed, the cells were washed once with 1x PBS, and cell lysates were harvested and RNA was extracted using acidic guanidinium thiocyanate-phenol-chloroform extraction [24]. The RNA was then DNAse treated and precipitated via phenol-chloroform extraction. Following precipitation, 1μg of RNA was reverse transcribed using random hexamer primer and qRT-PCR was carried out as described above with primer sets obtained from PrimerBank. The sequences of these primers may be found in the Supplemental Materials associated with this manuscript.

### Mouse Experiments

Four-week old C57BL/6 mice were obtained from Jackson Laboratory and were inoculated in the left, rear footpad with 1000 PFU of virus in diluent (1× PBS) in a volume of 10μL. Mock-infected animals were injected with diluent alone. Mice were monitored for neurological signs of disease and weighed twice daily. Neurological scoring is was as follows: 0 = No signs of overt disease, and normal behavioral activity; 1 = Abnormal trunk curl, grip, or tail weakness (1 of 3); 2 = Abnormal trunk curl, grip, or tail weakness (2 of 3); 3 = Absent trunk curl, lack of gripping, tail paralysis; 4 = Pronounced dragging of one or more limbs; 5 = hind or fore limb paralysis. On the termination day for each experiment or when mice met endpoint criteria (neurological scores of 5 or 4 if the animal was unable to obtain food or water), or weight loss greater than 20% of initial body weight, the mice were sedated with isoflurane and euthanized by thoracotomy. Blood was then collected and serum obtained by collecting blood in serum separator tubes. Following exsanguination, tissues were collected by dissection. Tissues were then placed in 1x PBS and homogenized using Kimble BioMasher II closed system micro tissue homogenizers. Ankle tissue was processed by bead beating using a Bead Ruptro 4 (Omni International). The infectious virus present in the tissue was quantified by plaque assay.

For histology, uninfected and SINV infected mice brains were removed at day 7 post infection and were divided in half sagitally. One half was used to asses viral titer (described above), while the remaining half was fixed in 4% formaldehyde and sectioned in paraffin. Tissue sections were then stained with haematoxylin and eosin (H&E). Pathological changes were scored by a board certified veterinary pathologist (through the Comparative Pathology Core Services facility, Iowa State University) in the indicated categories and regions of the brain were scored as follows: 0=normal, 1=minimal, 2=mild, 3=moderate, 4=severe.

### Detection of viral genome and cytokine transcripts in mouse brains

To measure the level of viral genome and cytokine transcripts in the brains of infected mice, RNA was extracted from brain homogenate using acidic guanidinium thiocyanate-phenol-chloroform extraction. The RNA was then DNAse treated and precipitated via phenol-chloroform extraction. Following precipitation, 1μg of RNA was reverse transcribed using Protoscript II reverse transcriptase (NEB) and random hexamer primer. The RNA genome was detected using BrightGreen Express qPCR master mix (abm good) and the following primer set specific for nsP1= F: 5-AAGGATCTCCGGACCGTA-3, and R: 5-AACATGAACTGGGTGGTGTCGAAG-3. A standard curve of known concentrations was used to determine the absolute quantities of viral genomic RNAs. Cytokine transcripts were detected using TaqMan Fast Advanced Master Mix and the Applied Biosystems TaqMan Array Mouse Immune Response plates (Catalog number:4414079) according to manufacturer’s instructions.

### Neuron Viability

Neuron viability was determined using a previously described method of ethidium bromide and acridine orange staining [25]. SK-N-BE(2) cells were seeded in a 96-well plate and, upon reaching 80-90% confluency, were inoculated with either wild-type SINV or one of the capping mutants at an MOI of 30 IU/cell. After a 1 h adsorption period, the inoculum was removed, the cells were washed once with 1x PBS, and DMEM/F12 media was added. At 24hpi, cell viability was assessed using ethidium bromide/ acridine orange staining described in Ribble et al. Briefly, the 96-well plate was centrifuged at 1,000 RPM for 5 min using an Allegra 25R model centrifuge (Beckman Coulter) with inserts for 96-well plates. Following centrifugation, 8μL of EB/AO dye solution (100μg/mL ethidium bromide and 100μg/mL acridine orange in 1x PBS) was added to each well. Cells were viewed using an epifluorescence microscope. Tests were done in triplicate and a minimum of 100 total cells per well were counted using ImageJ.

### Animal Ethics and Research

This study was carried out in strict accordance with the recommendations described in the Guide for the Care and Use of Laboratory Animals of the National Institutes of Health. The protocol was approved by the Institutional Animal Care and Use Committee of the University of Louisville (Approval #17-140). All manipulations which could result in acute pain or distress were performed under isoflurane anesthesia.

### Statistical Analysis

Unless otherwise stated, the quantitative data presented in this study represent the means of data from a minimum of three independent biological replicates. The in vivo studies described in this study were performed in duplicate using two independent preps of viral stocks. An area-under-the-curve approach was used to statically assess the growth curve data presented in figure 1 to determine the differences in viral growth kinetics through the course of the assay. Comparative samples were statistically analyzed as previously described [22], using variable bootstrapping where appropriate. The survival data presented in figure 4 were statistically analyzed using the log rank test. Student’s *t* test was used to determine the *P* values associated with individual quantitative data sets.

**Figure 1.**
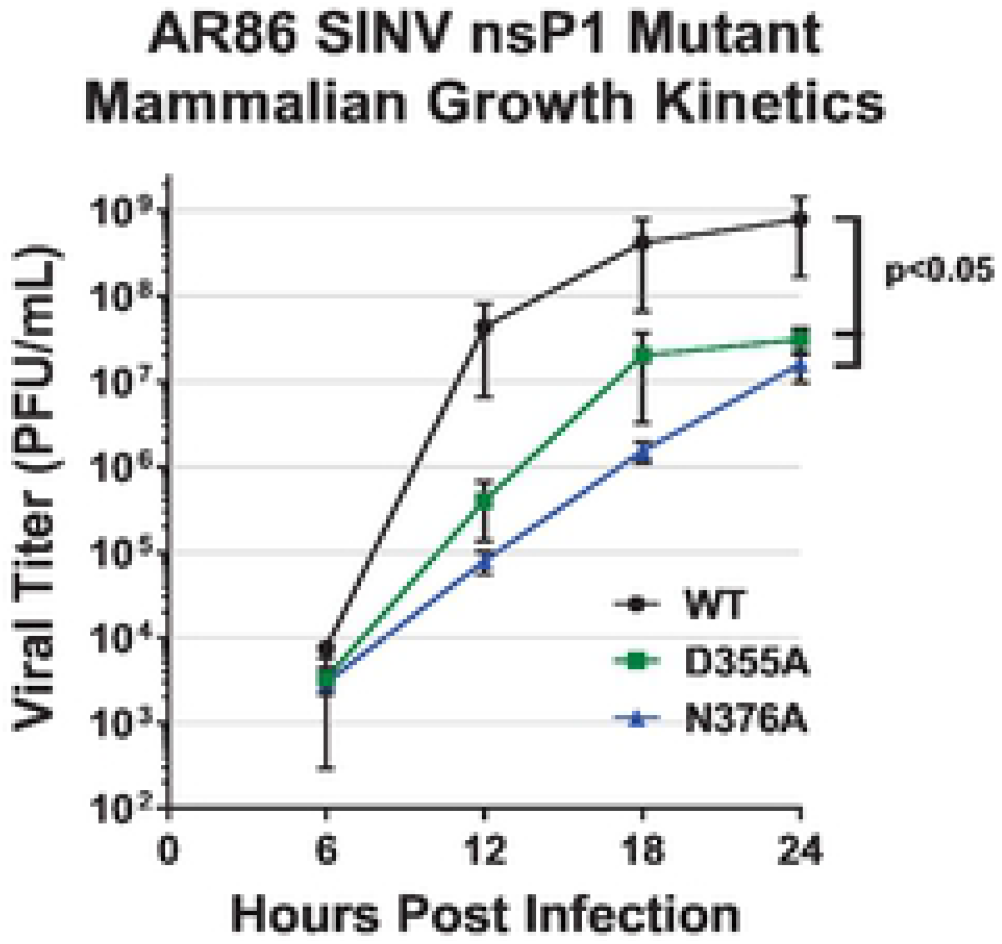
Altering viral capping efficiency negatively impacts infection of AR86 SINV in mammalian cells. One-step growth kinetics of the individual capping mutants and the parental wild-type SINV in BHK-21 cells infected at an MOI of 5 PFU/cell. All the quantitative data shown represent means of results from three independent biological replicates, with error bars representing standard deviation of the means. Statistical significance was determined by analysis of the area under the curve.

## RESULTS

### Altering Capping Efficiency Is Detrimental To Viral Growth Kinetics In Neurovirulent SINV *In Vitro*

Given our previously reported significant findings describing the molecular phenotypes of the nsP1 mutants in tissue culture models of infection, we were interested in characterizing how altering ncgRNA production impacted SINV infection *in vivo*. However, our previous characterizations of the capping mutants were done using a Toto1101-derived strain of SINV, which is tissue culture adapted and does not cause disease in adult wild type mice. Rather than rely on very young mice or mouse models with deficiencies in IFN competency to assess pathogenesis we elected to change the SINV strain used to allow assessments in adult wild type mice. Thus, we incorporated the D355A and N376A nsP1 mutations into the AR86 background of SINV, a neurovirulent strain capable of causing lethal disease in an adult mouse model. To confirm that these point mutations had similar phenotypes in the AR86 background as compared to the previously used Toto1101-derived background, viral growth kinetics were assessed in tissue culture models of infection (Figure 1). Similar to what we have reported previously, increasing the capping efficiency of nsP1 with the D355A mutation resulted in an approximately 1.5 log decrease in viral titer over the course of infection. Likewise, decreasing capping efficiency with the N376A mutation also resulted in a significant decrease in viral growth kinetics.

While the phenotype associated with the N376A mutant was more dramatic in the AR86 strain than what was reported in our previous study, this might be explained by the fact that the AR86 strain of SINV is not adapted for replication in tissue culture like the previously used Toto1101 strain. Therefore, the impact of the N376A mutation on replication is likely exacerbated in tissue culture systems, leading to the significantly reduced viral growth kinetics seen in Fig. 1.

The recapitulation of the original D355A phenotype and the pronounced N376A phenotype in the AR86 background provided a means by which the biological impact of the ncgRNAs on viral infection and pathogenesis could be assessed using an adult wild type mouse model. However, since our previous data suggested that the early events of the viral lifecycle may be altered by modulating capping efficiency, we decided to first characterize the engagement of the AR86-derived nsP1 mutants with the host innate immune response at a cellular level prior to utilizing a small animal model of SINV infection (LaPointe, 2018).

### Sensitivity To Type I IFN Correlates With Capping Efficiency In Tissue Culture

For the Old World alphaviruses, the translation of the alphaviral nonstructural proteins, specifically nsP2, is associated with the shut off of host transcription during SINV infection, and is subsequently also associated with viral resistance to type-I IFN [26]. We have previously shown that altering SINV capping efficiency correlated with changes to viral nonstructural protein expression early during infection [21]. Specifically, increasing capping efficiency with the nsP1 D355A point mutation lead to increased nonstructural protein expression, while the reverse was true when capping efficiency was decreased via the nsP1 N376A point mutation. Accordingly, we hypothesized that, due to increased expression of nsP2, the nsP1 D355A mutant would have decreased sensitivity to type-I IFN. Conversely, the nsP1 N376A mutant was hypothesized to have increased sensitivity to type I IFN, due to its decreased expression of nsP2.

To determine whether changes in viral genomic RNA translation affected the sensitivity of the nsP1 mutant viruses to exogenous type-I IFN, IFN-competent L929 cells were infected with either wild type or a SINV nsP1 mutant virus. Type-I IFN was then added at the indicated times post infection and viral titer was measured at 24hpi (Figures 2A-C) [27]. To comparatively determine how type-I IFN impacted viral infection of the capping mutants compared to wild type infection, the difference in viral titers between IFN treated and control infections were calculated and made relative to wild type titer differences (Fig. 2D). In wild type SINV infection, viral titer was reduced by ~2 logs when IFN was added at 0 hpi, but by 4 hpi, viral titer was equivalent to that of the no IFN control (Fig. 2A). As expected, the D355A mutant, which had increased capping efficiency and increased genomic translation, was found to be significantly more resistant to type-I IFN treatment early during infection compared to wild type SINV, with viral titer only being reduced by ~ 1 log when IFN was added at 0 hpi (Fig. 2B). However, when IFN is added 2hpi or later, the D355A mutant and wild type SINV were equally interferon resistant, as both viruses exhibited similar relative viral titers (Figure 2A, B, and D). When IFN is added at 4hpi, D355A viral titer is comparable to the interferon negative control and the difference in titer seen between the D355A mutant and wild type SINV is comparable to what was seen in figure 1A. In contrast, the N376A mutant, which had decreased capping efficiency and decreased nonstructural gene expression early during infection, was found to be have wild-type equivalent sensitivity to IFN when added at 0 and 1 hpi, but the SINV N376A mutant was significantly more sensitive when IFN was added at 2 hpi or later (Fig. 2C). Surprisingly, even when IFN was added as late as 4hpi, the titer of the N376A mutant remained decreased compared to the IFN negative control, revealing that by decreasing capping efficiency, the N376A mutant remains sensitive to IFN for a longer period of time than wild type SINV.

**Figure 2.**
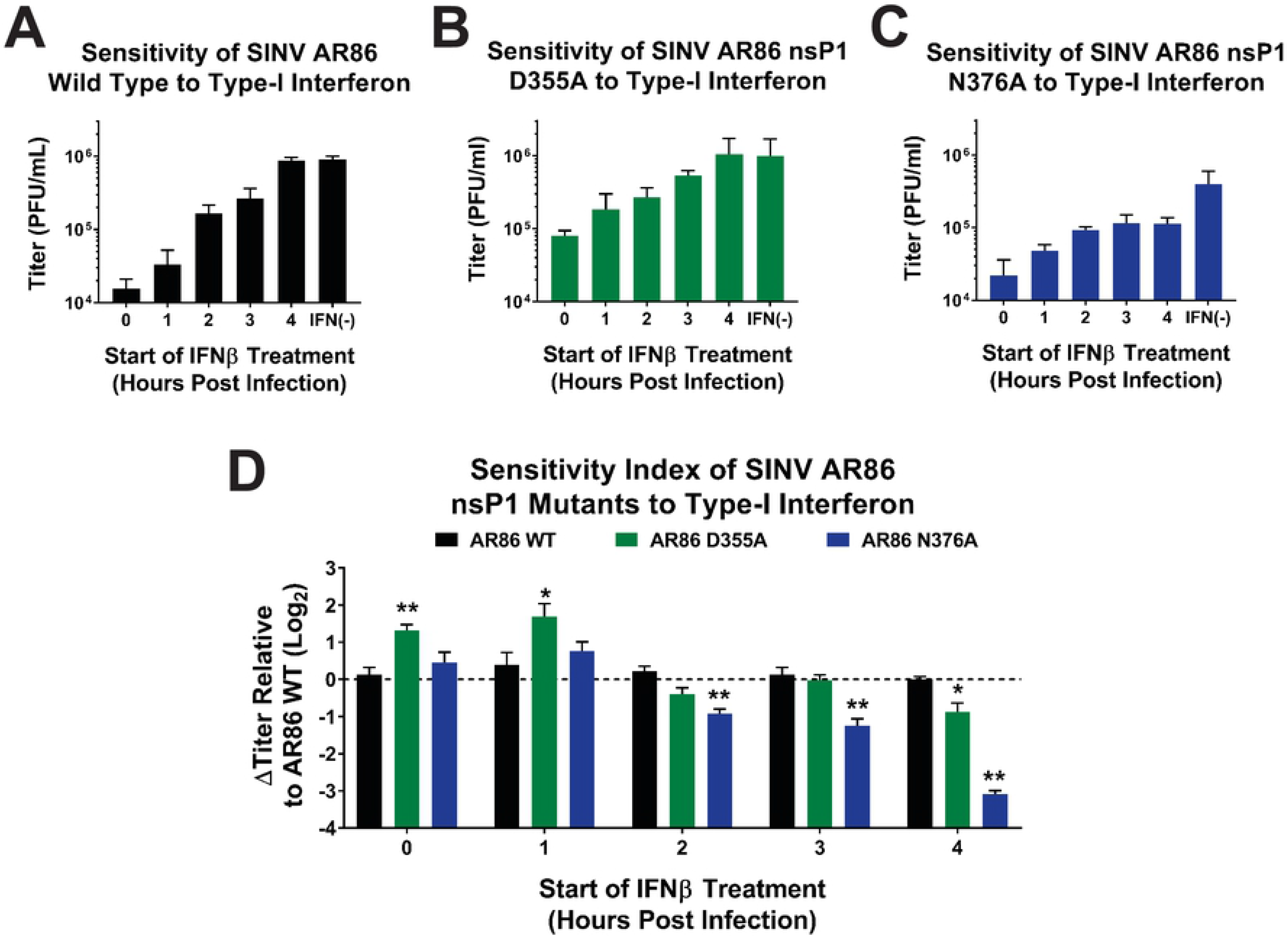
Analysis of SINV sensitivity to type-I interferon. L929 cells were infected with either wild-type SINV or an individual capping mutant at an MOI of 10 PFU/cell. At the indicated times post infection, 20 IU of recombinant Type-I IFN was added to the growth medium and the cells were incubated for a period of 24 hours. (A-C) Viral titers were quantified via plaque assay, and (D) the relative sensitivity of the viruses was determined by comparing their growth relative to untreated controls. All the quantitative data shown represent means of results from three independent biological replicates, with error bars representing standard deviation of the means. Statistical significance was determined by Student’s *t* test. *=p<0.05. **=p<0.01.

It is interesting to note that, in the absence of IFN treatment, infection of the L929 cells with wild type SINV or the D355A mutant resulted in roughly equivalent viral titers at 24hpi. This is different than what we observed previously in BHK-21 cells, where the D355A mutation resulted in significantly decreased viral titer (Fig. 1). Additionally, while the N376A mutant’s titer is still decreased compared to wild type virus in figures 2A and C, it is to a lesser extent than what was seen with BHK-21 cells in Fig. 1. This difference in phenotypes between cell lines is likely due to the fact that the L929 cells are IFN competent and will produce IFN in response to viral infection while BHK cells are incapable of doing so, resulting in differential viral replication rates due to the apparent differences in IFN sensitivity.

Overall, the sensitivity of the SINV nsP1 mutants to IFN reflects the differences seen in capping efficiency. Increased capping efficiency, and therefore increased genomic translation, resulted in the D355A mutant being more resistant to type-I IFN early during infection when compared to wild type SINV. Likewise, decreased capping efficiency, and hence decreased genomic translation early during infection, resulted in the N376A mutant being more sensitive to type-I IFN. This indicates that the viral response to type-I IFN, and its ability to mitigate the effects of IFN expression on viral replication are altered depending on the level of ncgRNA produced during infection.

### Modulating ncgRNA Production Alters The Host Type-I Interferon Response To SINV Infection

The capacity of the host cell to detect viral infection and mount an antiviral response by upregulating interferon and interferon stimulated gene (ISG) expression is an important aspect of viral infection. Given that altering capping efficiency significantly impacted the virus’ sensitivity to type-I IFN, we were interested in determining whether changes in viral capping efficiency would also impact the stimulation of the host type-I IFN response. To assess the extent to which the nsP1 mutants elicited an interferon response, interferon competent L929 cells were infected with either wild type SINV or one of the capping mutants and then the expression of IFNβ and several representative ISGs were measured at the transcriptional level. For this experiment, we intentionally selected ISGs with well-established times of maximal expression post viral infection, so that we could accurately determine the host antiviral response over the course of infection [28]. As such, SHB and IFIT2 were used to determine the ISG expression at 2 and 4hpi, CXCL10 and IFIH1 at 6 and 8hpi, Viperin and MX2 at 16hpi, and OAS2, BST-2, and IFNβ at 24hpi (Figure 3).

**Figure 3.**
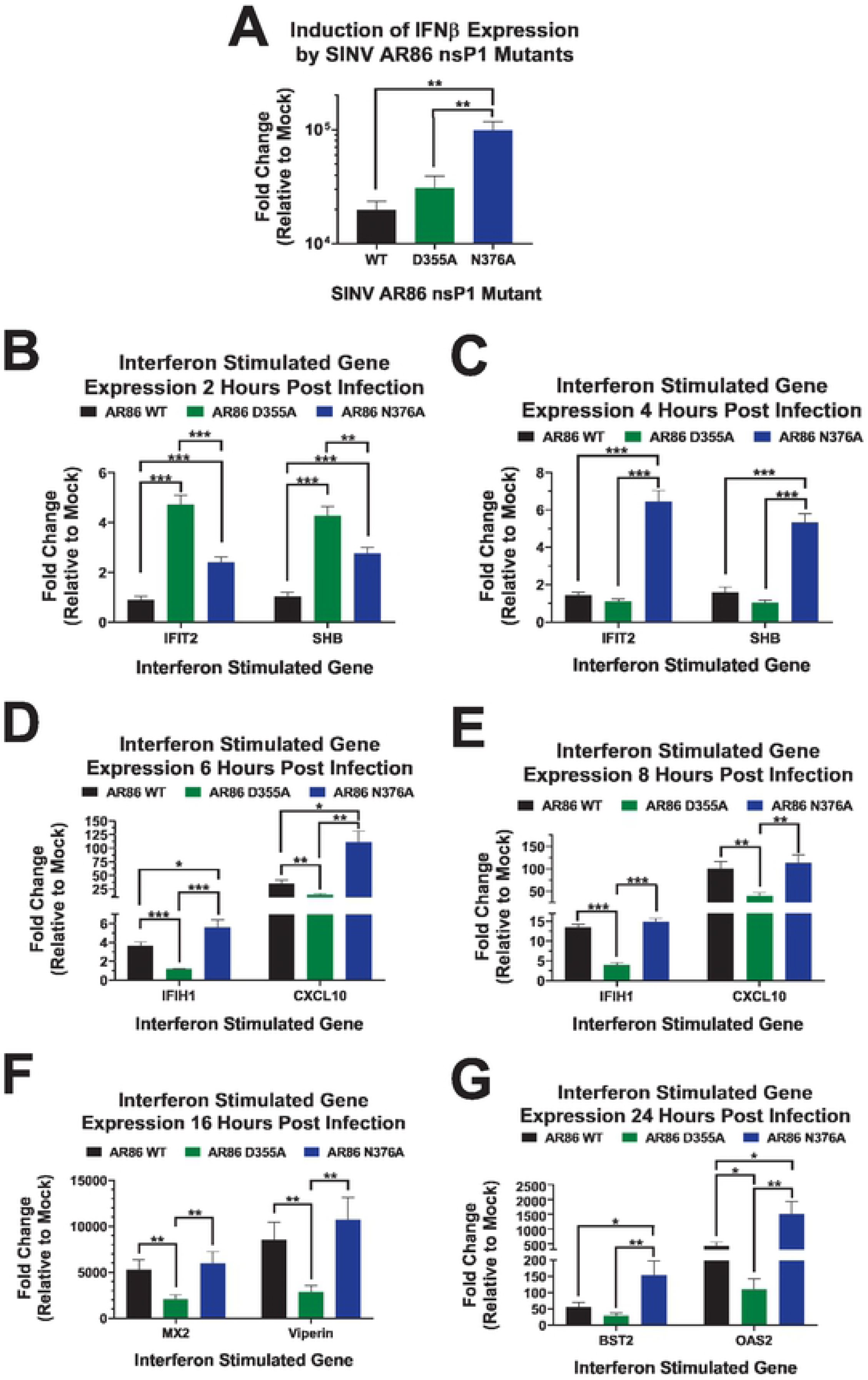
Production of type-I interferon and ISGs in response to SINV nsP1 capping mutants. L929 cells were infected at an MOI of 10 PFU/Cell with either wild-type SINV or an individual capping mutant. Cell lysates were collected at 2 (B), 4 (C), 6 (D), 8 (E), 16 (F), or 24hpi (A,G) and transcript expression levels were determined by qRT-PCR. Data was normalized to GAPDH and nsP1 and calculated relative to uninfected controls. All the quantitative data shown represent means of results from three independent biological replicates, with error bars representing standard deviation of the means. Statistical significance was determined by Student’s *t* test.

**Figure 4.**
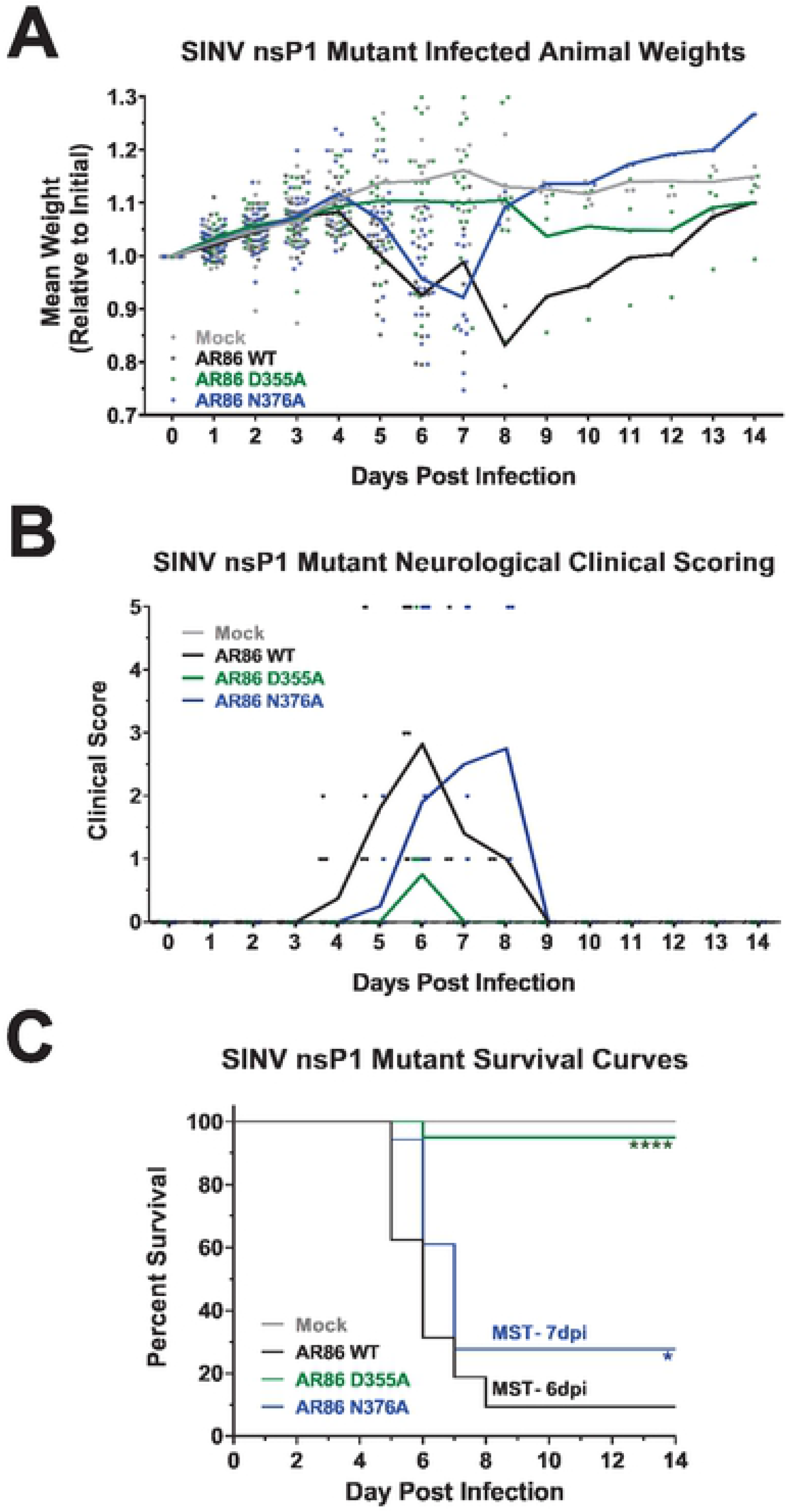
Increased vRNA capping efficiency reduces SINV AR86 mortality and pathogenesis. 4-week old male and female C57BL/6J mice were either mock infected or infected with 1,000 PFU of SINV AR86 wild-type, D355A, or N376A via rear footpad subcutaneous inoculation. Each data point represents a single animal from either experimental replicates. The experimentally infected mice were assessed over a 14 day period. (A) Animals were weighed twice daily. Weights are shown relative to initial weight after being infected. (B) Mice were scored based on a 1-5 scale for neurological response. (C) Kaplan-Meier analysis indicates the WT median survival time (MST) at ~6.4 days and the N376A mutant MST at ~7 days. The P values indicated on the figure were determined by the Log-rank test. *=p<0.05, ****=p<0.0001. Data shown were pooled from 2 independent experiments.

IFNβ expression at 24hpi was found to be similar between D355A and wild type infection, while N376A infection resulted in significantly greater IFNβ expression compared to both wild type and the D355A mutant (Fig. 3A). Looking at ISG transcript levels, both the D355A and N376A capping mutants showed increased IFIT2 and SHB expression at 2hpi as opposed to wild type SINV, which did not elicit a significantly increased ISG response compared to mock infection until 6hpi (Figure 3B). Infection with the N376A mutant continued to result in significantly increased ISG expression compared to wild type SINV until 8hpi, where transcript abundance became roughly equivalent to wild type infection (Figure 3C-E). However, by 24hpi, ISG expression once again were significantly increased (Figure 3G). In contrast, ISG expression in response to the D355A mutant was significantly decreased compared to wild type SINV by 6hpi, and remained so until 24hpi, where OAS2 expression was still significantly decreased, but BST2 expression was similar to wild type.

Overall, infection with the increased capping mutant, D355A, lead to a mostly decreased host antiviral response compared to what was seen during wild type infection, while the decreased capping mutant, N376A, elicited a response that was mostly increased compared to wild type SINV. The fact that the ISGs chosen within each timepoint had similar trends in expression further supports them being representative of the host antiviral response. Taken together, these results illustrate that modulating ncgRNA production has a significant impact on the host type-I IFN response.

### Increasing Capping Efficiency Significantly Attenuates Neurotropic SINV In A Mouse Model

As the capacity to avoid the elicitation of the host innate immune response, as well as shutting down the host type-I IFN response are vital to alphaviral replication and pathogenesis, the above data suggested that altering ncgRNA production could have profound effects on viral replication and pathogenesis *in vivo*. We hypothesized that, due to the nsP1 D355A mutant’s increased resistance to type-I IFN and generally reduced activation of ISG expression, mice infected with the nsP1 D355A mutant would experience disease severity similar to wild-type SINV infection, perhaps with the mean survival time being potentially decreased due to increased IFN resistance. Conversely, we hypothesized that mice infected with the N376A mutant would experience more mild disease and decreased mortality because of the mutant’s increased sensitivity to IFN and the greater expression of ISGs in response to infection in tissue culture models of infection. To test our hypothesis, we infected 4-week old male and female C57BL/6 mice with 1,000 PFU of SINV AR86 wild-type, nsP1 D355A, or nsP1 N376A via rear footpad subcutaneous inoculation. Mock infected mice were inoculated with PBS in the same manner. When infected with wild type SINV, adult C57BL/6 mice displayed significant weight loss as well as severe neurological symptoms, including rapid onset paralysis of the limbs, blindness, and seizures at approximately day 6 post infection (Figure 4A and B). Infection with wild type SINV also led to significant mortality, with infected mice having a mean survival time of ~6 days post infection (Figure 4C). Likewise, mice experimentally infected with the decreased capping mutant N376A exhibited similar weight loss and neurological symptoms as wild type infected mice. However, the onset of disease in the N376A infected mice was delayed compared to wild-type SINV, with neurological symptoms starting at 5 dpi with the mean survival time being ~7 dpi, a full day later than what was seen with wild type SINV. In addition, a slightly greater proportion of mice survived when infected with the N376A mutant as opposed to wild type SINV. This increase in survival may be due to the delay in the N376A mutant to cause neurological symptoms, allowing the mice to be slightly older, and therefore better able to resist severe, lethal encephalitis [29–31].

Surprisingly, the increased capping mutant virus D355A was significantly attenuated in mice. Compared to the previous two viruses, mice infected with the D355A mutant experienced minimal weight loss, milder neurological symptoms, and significantly reduced mortality, with all but one mouse surviving to the end of the study. Given that these mice did in fact show mild neurological and non-neurological symptoms and reduced weight gain compared to mock infected mice, we concluded that the nsP1 D355A increased capping mutant virus is indeed capable of causing pathogenesis, although the severity of disease is significantly reduced compared to wild type infection.

The trends seen in morbidity and mortality between the mice infected with wild type SINV versus the capping mutants were further reflected in H&E stained sections of the brains of infected and uninfected mice (Figure 5). The brain sections of both wild type and N376A infected mice displayed numerous lesions consisting of lymphocytic meningitis; perivascular cuffing, which is indicative of immune cell infiltration; and neuronal apoptosis, which left open pockets in the tissue (Figure 5A). In addition, mice infected with either wild type SINV or the N376A mutant had significant pathology in terms of inflammation, neuronal degradation, and glial cell proliferation in multiple areas of the brain. Pathology was highest in the cerebrum and the midbrain / brainstem, but lesions were also found in the hippocampus and medulla oblongata of some mice (Figure 5B). Conversely, the brains of the D355A infected mice resembled mock infected mice, with no immune infiltration, cell death, or other signs of pathology in any area of the brain. Given that viral killing of neurons is the speculated cause of encephalitis and paralysis in SINV infection, the differences in tissue damage and neuron death seen between the D355A mutant and the other two viruses were not unexpected, as the D355A infected mice did not exhibit signs of encephalitis or limb paralysis (Figure 4B) [32, 33].

**Figure 5.**
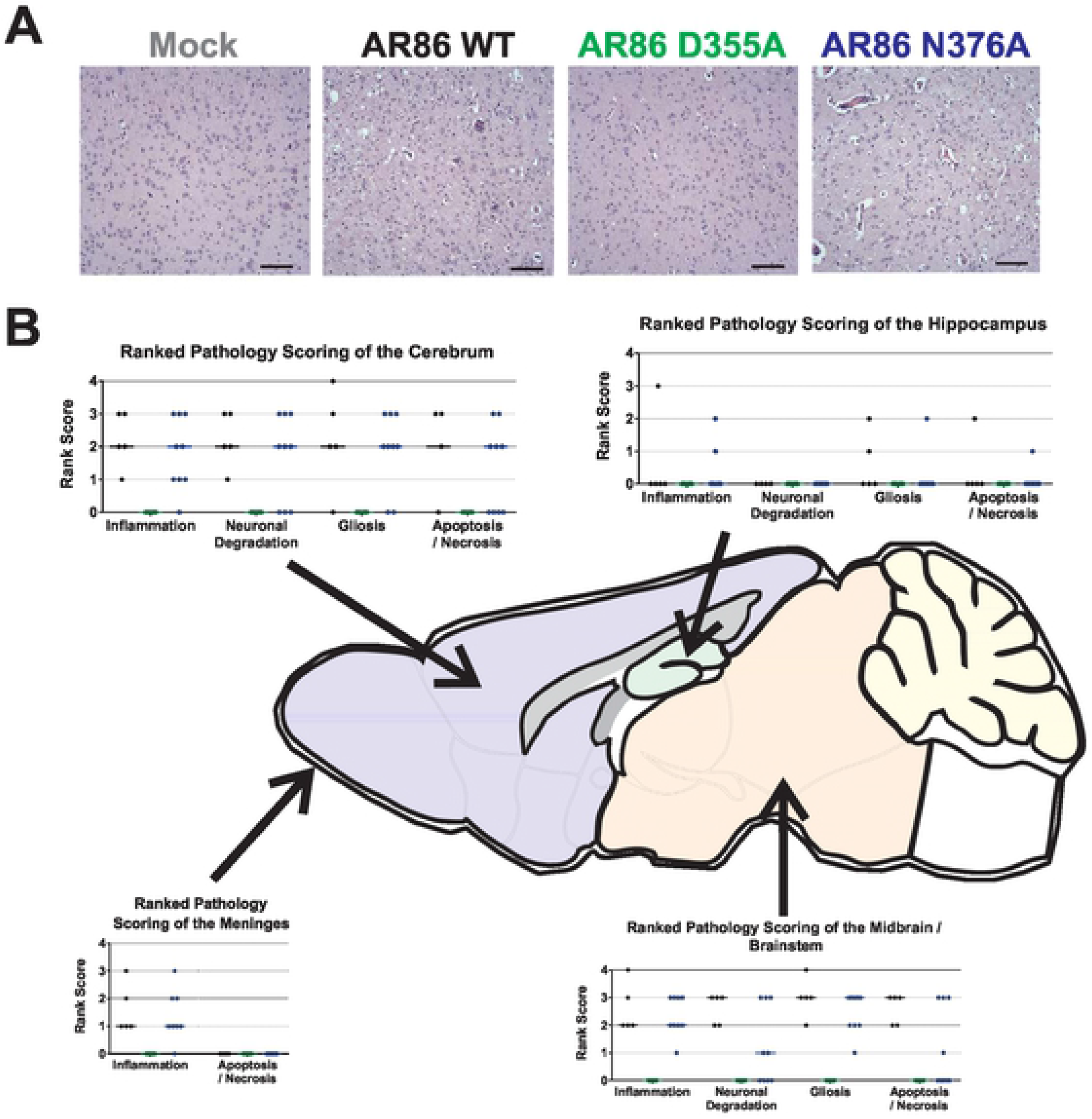
Increased capping efficiency leads to decreased pathology in the brain. A) Representative H&E stained sagittal sections of the midbrain (20× magnification) from mock, wild-type, or capping mutant infected mice at 7dpi or at the time at which end point criteria was met. The brains of SINV wild-type and N376A infected mice show large amounts of perivascular cuffing, immune infiltration, and cell death not present in the mock and D355A infected mice. Scale bar = 0.1mm. B) Ranked pathology scoring of indicated sections of the brain from infected mice. Data points indicate scoring for each experimental animal, representing at least 5 biological replicates.

Collectively, these data suggest that modulating the production of ncgRNA has significant impacts on alphaviral pathogenesis. Overall, the nsP1 N376A point mutation, which increased ncgRNA production through decreased capping efficiency, resulted in delayed disease progression and mortality compared to wild type SINV *in vivo*. However, severity of neurological symptoms and pathology in the brain were unaffected. In contrast, the nsP1 D355A point mutation, which decreased ncgRNA production through increased capping efficiency, resulted in significantly decreased mortality, mild neurological symptoms, and little to no pathology in the brain. While the presence of mild symptoms and a lack of weight gain in the D355A infected mice do suggest that the virus was capable of trafficking to the brain and replicating, these data do not eliminate these as being possible reasons for the decreases in pathogenesis seen thus far.

### Attenuation Of Viral Pathogenesis Is Not Due To Deficits In Viral Dissemination Or Replication

Given that the D355A and N376A nsP1 mutants showed decreased viral titers in tissue culture model systems compared to wild type SINV (Figure 1), we hypothesized that the reduction in mortality seen in figure 4 was due to poor viral replication, dissemination, or a change in virus tropism for the brain. In order determine the impact of modulating capping efficiency on viral replication and to confirm that the nsP1 D355A mutant did in fact make it to the brains of infected mice, we measured viral titer at the site of inoculation and in the serum at 1dpi as well as in the brain at 7dpi. By comparing the viral titers of the nsP1 D355A mutant to wild type SINV in these tissues, we will be able to determine if decreasing ncgRNA production impacts viral pathogenesis by altering viral replication, dissemination, or tropism to the brain. If the D355A mutant had defective dissemination or tropism to the brain, then we would expect to see wild type titers at the site of inoculation, as well as potentially the serum, but an absence of viral titer in the brain. Alternatively, if viral titers for the D355A mutant are significantly decreased in the ankle, serum, and brain, then this would suggest that the decreases seen in pathogenesis were due to poor viral replication and dissemination.

Surprisingly, in contrast to what was expected given our tissue culture data, viral titers in the ankle, serum, and brain were more or less equivalent between wild type SINV and the two nsP1 mutants (Figure 6). The similar titers found in the ankle between the nsP1 mutants and wild type SINV show that the reduced pathogenicity of the nsP1 D355A mutant is not due to a defect in viral replication at the site of inoculation (Fig. 6A). Likewise, since the nsP1 D355A mutant had titers equivalent to wild type levels in the serum, we can also conclude that viral dissemination was not negatively impacted (Fig. 6B). Finally, while the viral titer of the D355A and N376A mutant were both slightly decreased in the brain compared to wild type SINV, the lack of a significant difference indicates that increasing capping efficiency did not alter viral tropism to the CNS and that there is no overt defect in viral replication (Fig. 6C). Interestingly, while these results are different from what was previously observed during infection of BHK cells (Figure 1), the trends seen in the serum and brain, where the N376A mutant has slightly decreased viral titer compared to wild type SINV and the D355A mutant, was similar to what was seen with the L929 cells (Figure 2).

**Figure 6.**
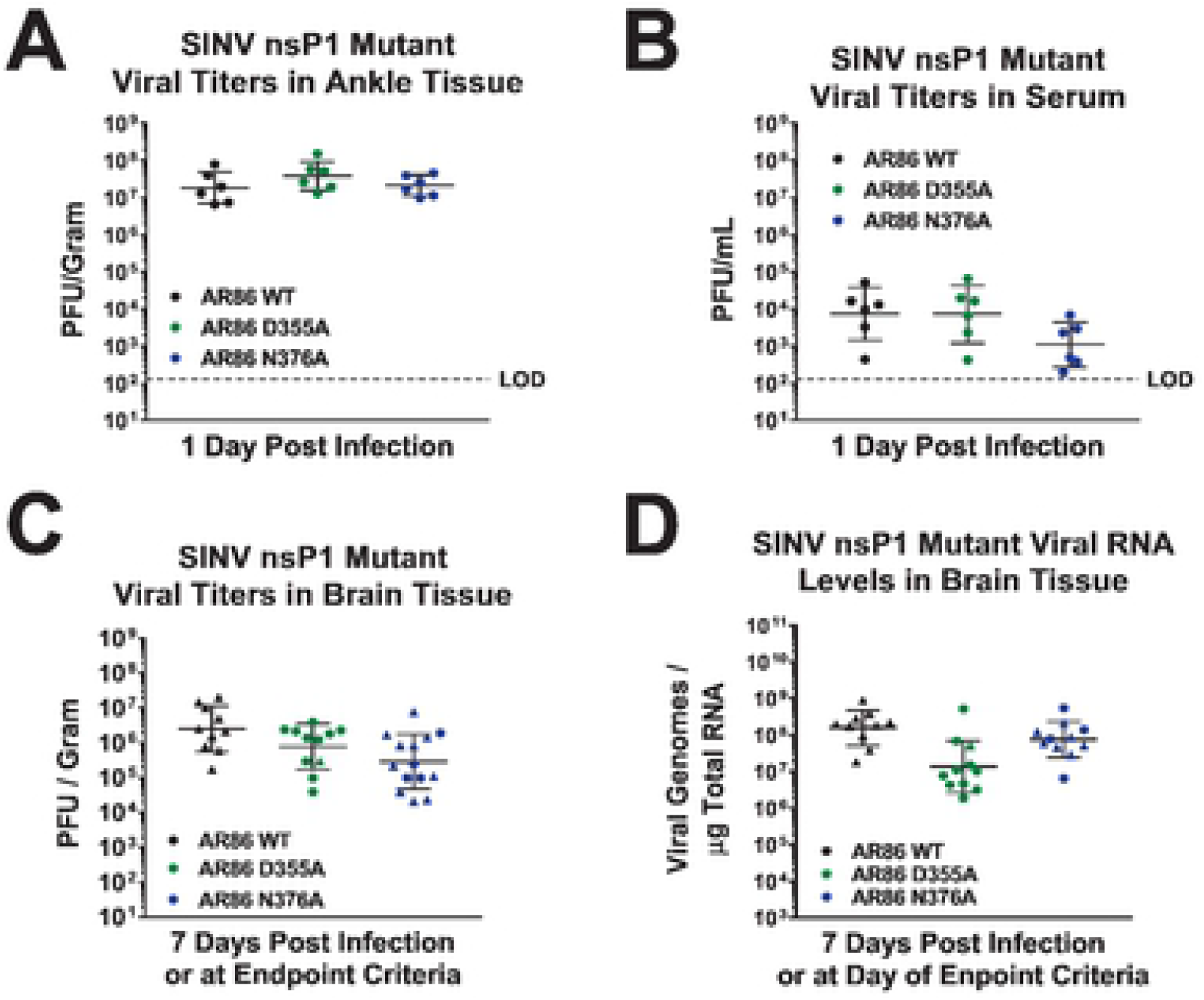
Viral replication is largely unaffected by altered capping *in vivo*. (A-C) Tissues were harvested at the indicated times post infection and viral titer was determined via plaque assay. (D) Viral genomes were measured by qRT-PCR. The data points indicate the individual titers for each experimental animal, and the mean values shown are the geometric means of at least four biological replicates from two independent experiments, with the error bars representing the geometric standard deviations of the means. ▲= mice that met endpoint criteria prior to day 7. Statistical significance was determined using Student’s t test.

The dissemination of the SINV mutants to the brain was further confirmed when viral RNA levels in the brain were measured (Figure 6D). While the nsP1 D355A mutant did exhibit slightly lower vRNA abundance in the brain compared to wild type SINV, it was not found to be a statistically significant difference and is likely an artifact due to differences in when the brain tissue was collected. While the majority of the D355A infected mice survived to 7 dpi when the brain tissue was collected, all of the wild type infected mice met endpoint criteria prior to 7dpi. As such, the adaptive immune response in the D355A infected mice that survived to 7 dpi may have started to clear some of the infected cells serving as viral RNA reservoirs from the brain, resulting in the decreased vRNA abundance compared to the wild type infected mice that did not survive long enough to mount a similar response. N376A RNA levels in the brain were also found to be equivalent to that of wild type SINV. These results indicate that, while the D355A and N376A nsP1 point mutations were capable of reducing viral titer in tissue culture, they were not detrimental to viral replication in mouse models of infection.

Taken together, these data suggest that both the increased capping virus D355A and the decreased capping virus N376A were capable of trafficking to the brain from the site of inoculation, and were capable of replicating to high titers within the brains of experimentally infected adult mice. It is also interesting to note that neither viral titer nor viral genomic RNA abundance correlated with death, as there were multiple mice which survived infection that had greater viral titer and vRNA abundance in the brain than mice which died. Overall, these results led us to conclude that the attenuation of pathogenesis seen in the D355A mutant were not due to either deficits in viral replication or tropism.

### Increasing Capping Efficiency Does Not Affect Viral Induction Of Neuronal Apoptosis

Since we did not find any significant differences in viral replication or tissue tropism / dissemination, we next determined if decreased mortality in the D355A mutant was due to an altered capacity to induce neuronal death, as virally induced apoptosis of neurons in the brain, brainstem, and spinal cord have been shown to be responsible for the severe neurological symptoms that arise during SINV infection [32–35]. In addition, our previous study characterizing the nsp1 capping mutants in tissue culture showed that both the D355A and N376A mutant demonstrated increased cell viability in BHK cells compared to wild type SINV, supporting the possibility that the D355A mutant might have differences in cell viability in neurons as well (LaPointe, 2018). In light of our previous study as well as the striking difference in cell death between the D355A and wild type infected brains (Figure 5), we hypothesized that increasing capping efficiency would decrease the virus’ capacity to kill neurons. To test this hypothesis, we infected SK-N-BE(2) cells with either wild type SINV or one of the capping mutants and determined cell viability 24 hours after infection using ethidium bromide and acridine orange staining. Infection with the decreased capping N376A mutant resulted in significantly greater cell viability compared to either wild type SINV or the D355A mutant, which both exhibited roughly similar levels of cell death (Figure 7). Interestingly, these results suggest that increasing ncgRNA production leads to increased cell survival in tissue culture models of infection, while decreasing ncgRNA production does not seem to impact the virus’ capacity to kill neurons in tissue culture models of infection. Given this, we can conclude that the differences seen in tissue damage between the D355A and wild-type SINV infections in figure 5 are not solely due to deficits in the D355A mutant’s capacity to kill neurons. Instead, the above results suggest that the neuronal death seen during SINV infection in mice may be largely due to the host antiviral inflammatory response rather than direct cell death due to infection.

**Figure 7.**
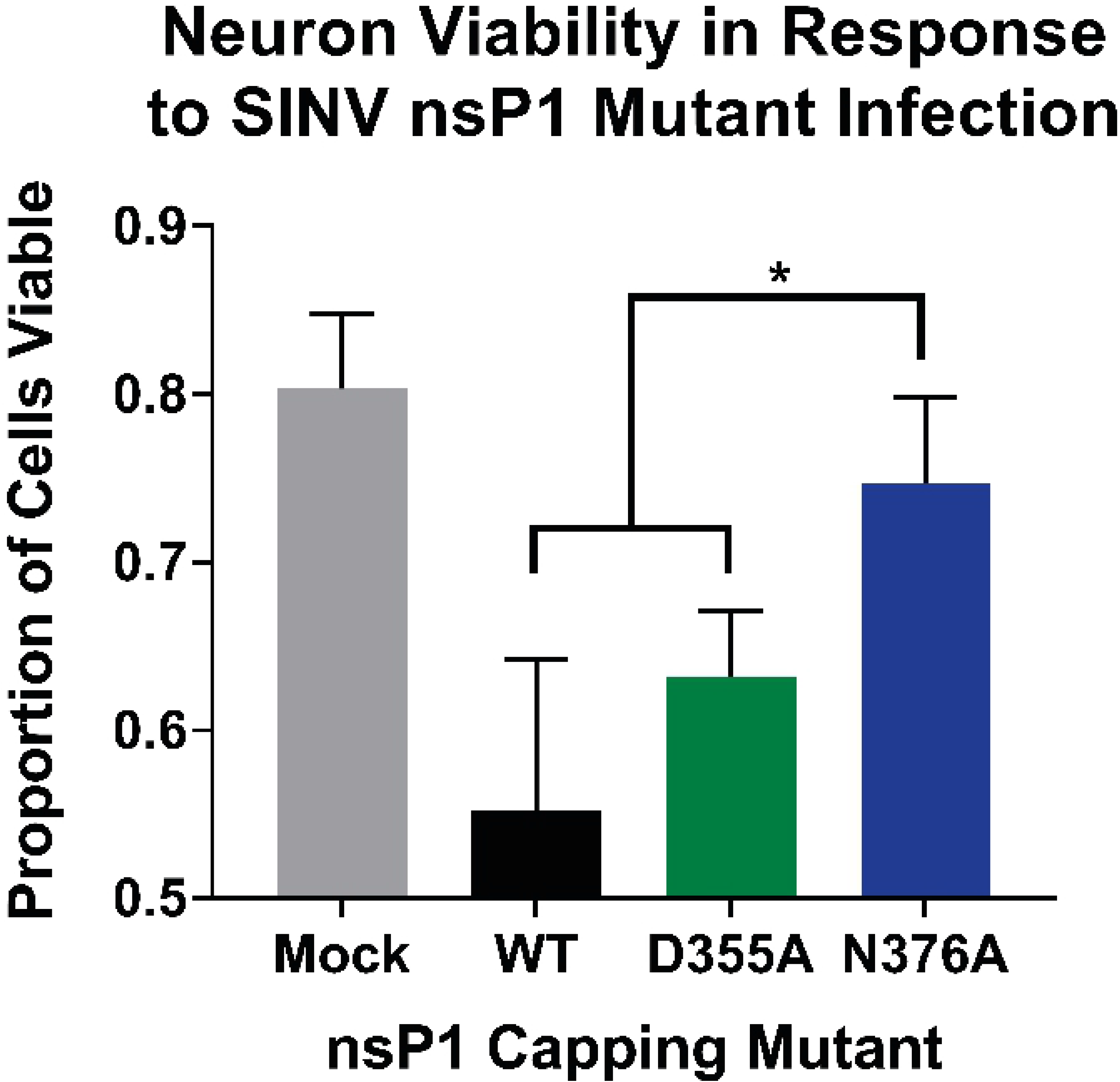
Neuron viability increased with decreased capping efficiency. SK-N-BE(2) neurons were infected at an MOI of 30 PFU/Cell with either wild-type SINV or an individual capping mutant. Cell viability was determined at 24hpi using ethidium bromide/ acridine orange staining and is represented as the proportion of viable cells out of total cells counted. A minimum of 100 total cells per well were counted using ImageJ. All the quantitative data shown represent means of results from three independent biological replicates, with error bars representing standard deviation of the means. Statistical significance was determined by Student’s t test.

### Differential ncgRNA Expression Alters The Immune Response To Infection

Because the differences in morbidity and mortality between the D355A nsP1 mutant and wild type SINV infected mice could not be explained by reductions of viral titer or the capacity to induce neuronal death, we next questioned whether differences in pathogenesis could be due to an altered host immune response. Given the reduced immune infiltration and inflammation seen in the D355A infected mice compared to the wild type or N376A infected mice (Figure 5), we expected infection with the D355A mutant to also result in the decreased expression of pro-inflammatory genes. To survey the immune response to SINV nsp1 mutant virus infections, RNA was isolated from whole brain homogenates of infected mice at 7 dpi or upon meeting endpoint criteria, and the level of select cytokine and chemokine transcripts were measured via a qRT-PCR based array. Out of the transcripts measured, 50 were found to be significantly increased in wild type infected versus mock infected mice (Figure 8A). When wild type and D355A infections were compared, we found 15 inflammatory cytokines and chemokines whose expression was determined to be significantly decreased by a magnitude greater than 2-fold (Figure 8B). These included chemokines involved in recruiting innate immune cells, such as CCL2, CCL3, and CXCL10, as well as important drivers of inflammation like IFNβ, IL1β, TNFα. In particular, the expression of CCL2 and CCL3 have been highly correlated with areas of the brain experiencing high levels of gliosis and apoptosis, which is consistent with the differences in pathology scoring in those areas between the D355A mutant and wild type SINV (Figure 5B) [36]. Interestingly, several of the proteins that had decreased expression in the D355A infected mice compared to both N376A and wild type infected mice where identified by gene ontology as being part of the extrinsic apoptotic pathway [37, 38]. The identification of proteins involved in apoptosis, specifically Fas, Il1α, Il1β, and TNFα, is consistent with the significant decrease in neuronal apoptosis seen in the H&E stained sections of mice infected with the D355A mutant. Surprisingly, IFNγ expression, which has been previously shown to be important for noncytolytic clearance of virus from neuronal cells, was not found to be significantly different in either mutant compared to wild type (Figure 8A) [39]. As expected, the decreases seen in the expression of these chemokines and pro-inflammatory cytokines are consistent with the reduced immune infiltration and tissue damage seen in Figure 5.

**Figure 8.**
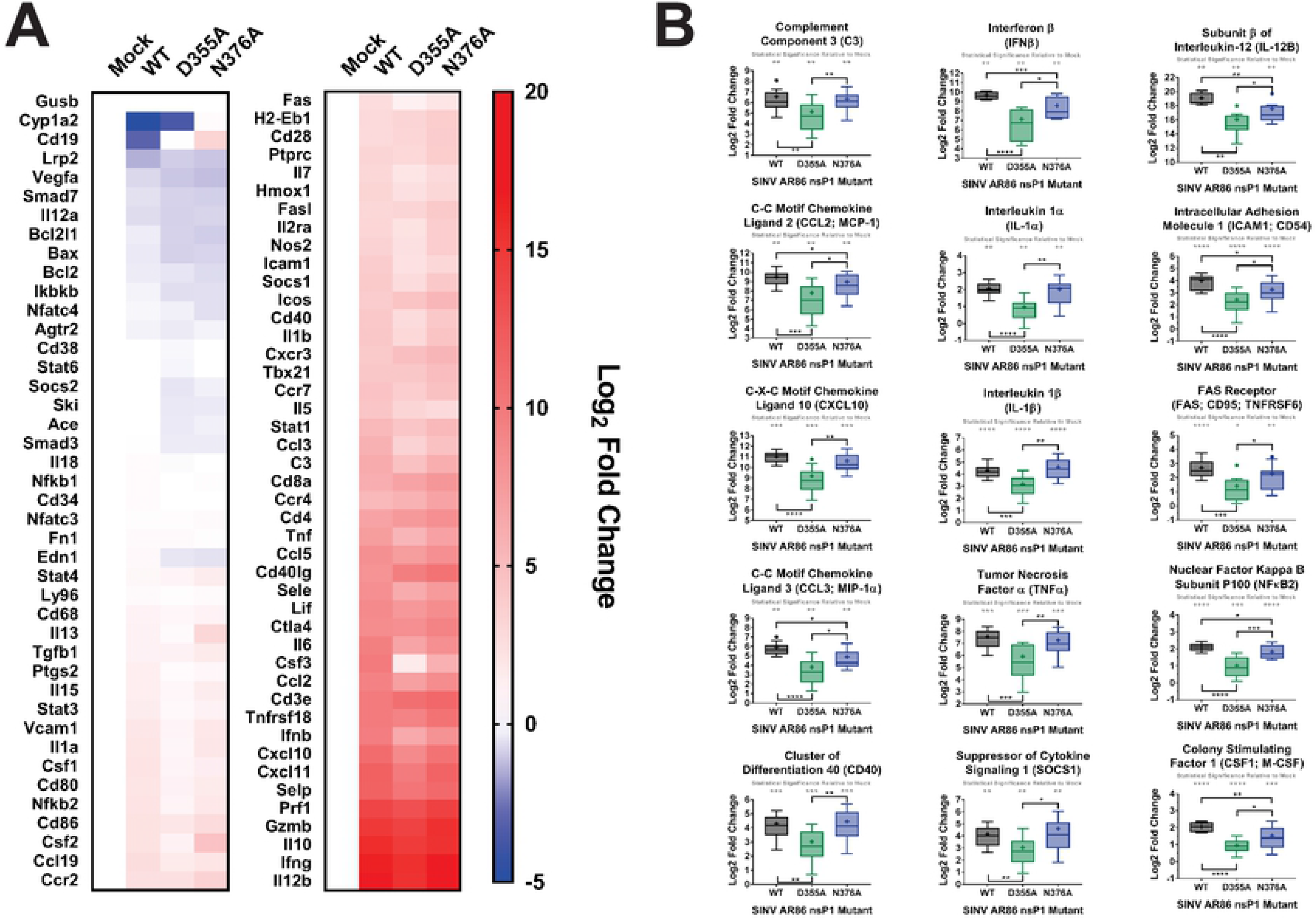
Increased viral capping efficiency results in reduced expression of pro-inflammatory genes in the brain. (A) Cytokine transcript levels in the brain at 7dpi were measured by qRT-PCR. Data was normalized to GAPDH and calculated relative to uninfected controls. (B) Cytokines and chemokines whose expression was significantly increased compared to uninfected controls and exhibited a significant difference in expression between wild type SINV and the D355A mutant that was greater than 2-fold. All the quantitative data shown represent means of results from at least three independent biological replicates, with center lines representing the median, + representing the mean, boxes representing the interquartile range, error bars representing standard deviation of the means, and • representing outliers as determined by Tukey’s method. Statistical significance was determined by Student’s t test.

Overall, these data show that the D355A increased capping mutant has significantly reduced pathogenicity in a wild type mouse model of infection correlating with decreased expression of multiple pro-inflammatory molecules. Furthermore, the above data suggests that it is likely the host response rather than viral replication per se that determines the extent of alphaviral pathogenesis. This is supported by the finding that the wild type, D355A, and N376A viruses all had roughly equivalent viral titers in the brain, yet wild type SINV and the N376A mutant have significantly increased pro-inflammatory cytokine expression compared to the D355A mutant. Taken together, it can be concluded that decreased ncgRNA production via increased capping efficiency leads to an altered host immune response, which in turn shapes alphaviral pathogenicity.

## DISCUSSION

### Altering Capping Efficiency Changes Viral Sensitivity To And Activation Of Host Type-I IFN

For Sindbis virus, the sensitivity of the virus to type-I IFN is largely dependent on the translation of the nonstructural proteins, especially nsP2, which are responsible for interfering with the IFN signaling pathway and for shutting down host transcription [26, 40–42]. Given our previously published work which showed that genomic translation correlated with nsP1 capping efficiency, it was not surprising that the increased capping D355A mutant showed increased resistance to type-I IFN while the decreased capping N376A mutant showed increased sensitivity to type-I IFN [21]. The fact that the D355A mutant showed increased resistance to type-I IFN even when IFN was added concurrently with the virus demonstrates how increasing capping efficiency allows the virus to more readily mitigate the effects of the IFN response. The D355A mutant’s increased resistance to IFN at such an early timepoint suggests that, in addition to increased translation, the mutation may also affect the timing of viral translation. Earlier translation of the viral proteins would allow the virus to shut down IFN signaling pathways, host PAMP sensors, as well as host transcription and translation earlier than the wild type virus, which would explain why the D355A mutant wasn’t as affected by IFN treatment [27].

In addition to changes in viral sensitivity to IFN, we also found that altering SINV capping efficiency resulted in changes in host antiviral gene expression during infection. Specifically, infection with the N376A mutant lead to a general increase in IFNβ as well as early and late ISG expression compared to wild type SINV. Meanwhile, infection with the D355A mutant lead to generally decreased ISG expression, with exception of the very early and very late time points which had either increased or wild type levels of expression. The altered IFNβ and ISG expression seen with the capping mutant likely reflects changes in the virus’ abilities to both avoid detection by the host as well as suppress the cellular antiviral response [40]. Because IFNβ and ISG transcript abundance was normalized to viral RNA levels, we know that the differences seen in antiviral transcript expression are not simply due to differences in viral replication or the amount of vRNA present. Rather, the changes observed in IFNβ and ISG expression for the D355A and N376A nsP1 mutants is likely due to both differences in PAMP production and their abilities to shutoff host transcription and translation. For example, both mutants elicit ISG expression earlier than wild type SINV, which is likely due to increased PAMP production. Specifically, the D355A mutant is known to have significantly increased genomic translation early during infection, which would result in accelerated / increased cell stress; while the nsP1 N376A mutant is known to produce more noncapped RNA, which has also shown to activate the host antiviral response [20, 21, 43–45]. For the D355A mutant, ISG levels decrease to wild type levels by 4hpi and then further decrease by 6hpi, at which time the increased expression of nsP2 is likely allowing the D355A mutant to more quickly shut off host transcription [27]. The equivalent levels of IFNβ and ISG transcripts seen 24hpi between the D355A mutant and wild type SINV may reflect when the wild type virus was able to reach similar levels of host shut off as the D355A mutant. Likewise, the sustained increase in ISG expression as well as increased IFNβ expression seen in the N376A mutant is likely a result of delayed and / or inefficient shut off of host transcription due to its decreased nsP2 translation. In addition, decreased viral translation may result in reduced viral interference with the JAK-STAT signaling pathway, leading to increased ISG expression [17, 27].

### Noncapped Genomic vRNAs Are Critical For SINV Pathogenesis In Mice

The ability to resist and shut down the type I IFN response has been shown to be one of the major determinants of virulence for SINV. For example, the AR86 strain of SINV is a virulent strain known to efficiently suppress the type I IFN response and cause lethal neurotropic disease in adult mice. The genetically similar Girdwood strain only partially inhibits the type I IFN response, and thus is avirulent in adult mice [17]. Given the above finding that the D355A mutant was both more resistant to type I IFN treatment and resulted in decreased ISG production compared to wild type virus, we were surprised to find that this mutant did not cause severe disease or mortality in mice. Equally surprising was the result that infection with the N376A mutant was similarly as severe and lethal as infection with wild type SINV, despite N376A being more sensitive to IFN treatment and stimulating more ISG expression *in vitro*. Taken together, this suggests that there may be a balance between inhibiting and activating the IFN response which results in pathogenesis, and that tipping the scales too far in either direction causes the virus to become avirulent. Furthermore, our results indicate that ncgRNA play a critical role in determining whether the virus is neurovirulent or avirulent, as decreasing ncgRNA production with the D355A mutation resulted in significant decreases in morbidity and mortality while increasing ncgRNA production with the N376A mutation resulted in fully neurovirulent virus and lethal disease.

Although the D355A mutation had a much more striking impact on morbidity and mortality, the N376A mutation also had noticeable impacts on alphaviral pathogenesis, namely a delay in the onset of symptoms, an increased mean survival time, and a moderate increase in overall survival compared to wild type SINV. The idea that inhibiting capping efficiency is detrimental to alphaviral infection is not novel and multiple compounds and drugs have been developed to specifically target nsP1 capping activity [12–15]. Our results do not negate the idea that decreasing or inhibiting capping efficiency is an effective means of combating alphaviral infection, but rather suggest that there is a threshold which needs to be reached before decreasing capping efficiency will be significantly detrimental to viral pathogenesis. Likewise, our mortality studies imply that the virus is much more sensitive to increasing capping efficiency and that this novel approach may be more effective in limiting the severity of viral disease.

During SINV infection, the development of severe encephalitis which leads to paralysis and death is caused by the extensive apoptosis and necrosis of neurons in the brain and CNS, which is believed to be virally induced [32, 35, 46, 47]. While apoptosis is the fate of many infected cells and neurons, a significant portion of uninfected neurons are killed during viral infection due to glutamate excitotoxicity [48]. The lack of apoptosis and necrosis seen in the brains of mice infected with the D355A mutant (Fig. 5) suggests that, during *in vivo* infection, decreasing ncgRNA production by increasing nsP1 capping efficiency leads to decreased viral induction of neuronal apoptosis and therefore decreased glutamate release, resulting in the reduced death of infected and uninfected neurons. However, when tested in tissue culture cells, infection with the nsP1 D355A mutant resulted in wild type levels of neuronal death, while infection with the N376A mutant, which showed extensive signs of neuronal apoptosis and necrosis *in vivo*, resulted in significantly increased neuron survival. Since increasing capping efficiency did not seem to impact the virus’ ability to induce neuron death in tissue culture, these data suggest that the apoptosis and necrosis of neurons seen in the brain during SINV infection is largely mediated by an external force, such as by the host immune response. This is supported by previous studies which propose that the majority of neuronal death due to apoptosis and glutamate excitotoxicity may be the work of T cells and astrocytes, rather than being directly virus-induced [49, 50].

Another surprising result was that disease severity did not correlate with increased viral titer or vRNA abundance. This is most clearly seen in the N376A infected mice, where the viral titers and vRNA levels in the brains of mice that died are interspersed among those which survived. In addition, surviving mice from both the D355A and N376A infections had titers and vRNA levels in the brain that were roughly equivalent to those found in the mice infected with wild type SINV which died. Furthermore, neither viral titer nor vRNA burden in the brain correlated with levels of inflammation seen in Figure 5. This suggests that, during SINV infection, high viral titer alone is not sufficient to cause severe disease in mice. In addition, altering capping efficiency did not significantly impact viral replication, dissemination, or tropism. This was illustrated by the roughly wild type-equivalent titers found in the ankle, serum, and brain indicating that both of the capping mutants were able to efficiently replicate at the site of inoculation, disseminate into the blood, and traffic to the brain. However, the slight decrease in N376A titer seen in the blood and brain does potentially suggest that dissemination may be slightly delayed or impaired when capping efficiency is decreased, and may explain the slight increase seen in mean survival time.

Unfortunately, one question we were unable to answer in this study was whether capping efficiency in brain tissue is similarly affected by the nsP1 mutations, as was previously shown in tissue culture model systems and with recombinant proteins [21, 51]. Regrettably, the limitations in sensitivity of previously established and currently available assays render us unable to directly answer this question, as these methods require a significant quantity of high-quality viral RNA that is difficult to obtain from brain tissue. This is likely due to the fact that an exceptionally few number of cells in the brain are required to be infected for the manifestation of significant disease and the appearance of endpoint criteria. However, the altered pathogenesis seen in mice in the absence of any obvious defects in viral replication, dissemination, or tropism lead us to believe that the point mutations incorporated are still altering capping efficiency and are not significantly affecting nsP functions in other ways. Previously characterized mutations in nsP1 which resulted in loss of neurovirulence did so by significantly altering vRNA synthesis and/or processing of the nonstructural polyprotein, which typically resulted in decreased viral titer in animal models of infection [16, 17, 52, 53]. Given that neither the D355A nor N376A mutations significantly altered viral titer or vRNA burden during SINV infection, as well as our previous paper showing that vRNA synthesis and polyprotein processing where unaffected, we can conclude that the phenotypes seen both in tissue culture and *in vivo* are the result of the mutations altering ncgRNA production through modulating capping efficiency. Furthermore, given the conserved effect of the D355A and N376A nsP1 point mutations in multiple alphaviruses in tissue culture and *in vitro*, it is likely that these mutations still respectively increase or decrease capping efficiency *in vivo*, but the magnitude that capping efficiency is altered may be different from what was previously seen in tissue culture [21, 51].

### Noncapped Genomic RNAs Determine SINV Virulence by Modulating the Host Inflammatory Response

Infection with the D355A capping mutant resulted in the decreased expression of multiple cytokines and chemokines associated with the recruitment of immune cells, regulating inflammation, and apoptosis. While there were a small number of cytokines found to be differentially expressed during infection with the N376A mutant compared to sild type SINV, they did not implicate any pathways in particular. Interestingly, expression of anti-inflammatory molecules such as TGFβ and IL-10 was found to be similar between D355A and wild type infection, while others such as SOCS1 were found to be significantly decreased. This suggests that the decreased inflammation seen with the D355A mutant is due to decreased activation of antiviral and inflammatory pathways rather than increased expression of anti-inflammatory cytokines. The decreased activation of these antiviral pathways is are likely due to both the reduced release of DAMPS from dying cells and decreased sensing of viral PAMPS. The first is supported by the identification of several of the affected proteins being involved in apoptosis as well as the decreased level of cell death seen with the D355A mutant. The second is supported by the decreased ISG expression seen during D355A infection prior to when host shutoff occurs (Figure 3). The decreased sensing of viral PAMPS may be due to the D355A mutant either being more efficient at inhibiting the cell’s viral sensors and signaling pathways through shutoff of host transcription or could be due to the D355A mutant producing fewer noncapped RNA, which are an established PAMP [43, 44]. Given that viral infection in animals is a continuous process, shut-off of cellular transcription and suppression of the IFN response in tissues likely does not occur as efficiently or completely as it does in cell culture, where all the cells are infected simultaneously. Therefore, the decreased production of inflammatory cytokines seen with the D355A mutant is likely due to reduced detection of DAMPs and PAMPs caused by decreased cell death and decreased production of ncgRNA. How exactly the ncgRNA are sensed by the host during viral infection is not currently known and is an ongoing interest in the Sokoloski lab. While there is some evidence that suggests that the noncapped RNA produced during alphaviral infection are at least in part sensed by RIG-I, it is unknown if this is also true for ncgRNA and there may be additional methods for detecting noncapped vRNA that have yet to be characterized [44, 54]. Overall, the correlation between decreased inflammation and decreased ncgRNA production leads us to conclude that the ncgRNA play a critical role in determining the host response to viral infection.

In conclusion, we have identified a novel determinant of Sindbis virulence which operates through a separate mechanism than those previously described. Specifically, decreasing the production of ncgRNA by increasing capping efficiency results in the loss of neurovirulence which we believe is due to the reduced production of RNA PAMPs that would otherwise cause excess inflammation and wide-spread cell death in the brain. The D355A mutation differs from previously identified nsP1 virulence determinants in that it does not negatively affect viral titer nor resistance to IFN, such as is seen with the SINV nsP1 cleavage mutant T538I and the 6 nsP1 mutations characterized in Ross River virus [16, 19]. The D355 residue in nsP1 is also unique from the aforementioned mutation sites in that it is very highly conserved among SINV strains as well as across both the old and new world alphaviruses. While the results of this paper indicate that the production of ncgRNA is critical to SINV pathogenesis, more work is needed to further characterize the mechanisms by which ncgRNA contribute to alphaviral neurovirulence.

## ACKNOWLEDGEMENTS

We thank the members of the laboratories of Drs. KJ Sokoloski, D Chung, and IS for their valuable input during the development and execution of this project, and the preparation / editing of the manuscript.

This work was funded by grants from the National Institute of Allergy and Infectious Diseases (NIH-NIAID), specifically R01 AI153275 to KJS; and by a COBRE program grant from the National Institute of General Medical Sciences (NIGMS), P20 GM125504 to KJS and R. Lamont. ATL was supported by an NIH-NIAID funded predoctoral fellowship, T32 AI132146. Additional support was received from the Integrated Programs in Biomedical Sciences (IPIBS) to ATL, and a generous startup package from the University of Louisville to KJS.

This work was supported in part by a grant from the Jewish Heritage Fund for Excellence Research Enhancement Grant Program at the University of Louisville School of Medicine.

The funding agencies had no role in the study design, data collection and analysis, decision to publish, or the preparation of the manuscript.

